# Network measures from the REWIRED simulation framework enhance prediction of post-stroke aphasia severity

**DOI:** 10.64898/2026.05.19.726069

**Authors:** Isaac Falconer, Maria Varkanitsa, Emerson Kropp, Swathi Kiran

**Author notes:** **Corresponding Author:** Isaac Falconer, 111 Cummington Mall, Suite 280, Boston, MA 02215.

## Abstract

Predicting post-stroke aphasia severity remains challenging, in part because language outcomes reflect not only focal cortical damage but also widespread disruption of structural and functional networks. Computational models of large-scale cortical dynamics offer a principled way to infer these network-level consequences from patient-specific lesions. Here, we present and evaluate REWIRED, a lesion-informed cortical dynamics framework designed to simulate individualized alterations in large-scale brain network organization after stroke. We first evaluated whether simulation-derived functional connectivity captured patient-specific variation in empirical functional connectivity beyond lesion burden and structural disconnection alone. We then developed a multiscale feature set combining lesion volume, lesion distribution patterns, probabilistic disconnectome metrics, and simulation-derived measures of functional connectivity and effective information flow (EIF). Finally, using a nested support vector regression (SVR) framework in a separate dataset, we tested whether simulation-derived features improve prediction of chronic aphasia severity, measured by the Western Aphasia Battery - Revised Aphasia Quotient (WAB-AQ), beyond lesion-distribution and structural-connectivity predictors.

Simulation-derived functional connectivity significantly predicted empirical functional connectivity beyond local lesion burden and structural disconnection alone. With respect to WAB-AQ prediction, lesion-based (Set 1) and disconnectome-based (Set 2a) features alone yielded modest accuracy. Adding simulation-derived features (Set 2b) produced substantial gains, and the full feature set (Set 3) achieved the best performance (RMSE = 14.5; r = 0.83), reaching accuracy that is competitive with recent multimodal neuroimaging approaches, despite relying solely on lesion-distribution inputs. EIF measures were consistently selected as top predictors, indicating that disruptions in interregional communication patterns carry behaviorally relevant information not captured by structural features alone.

These results support REWIRED as a framework for linking structural injury to distributed network dysfunction and behavioral outcomes. By integrating lesion information with large-scale cortical dynamics modeling, REWIRED provides a foundation for future individualized modeling of recovery and rehabilitation.

## 1. Introduction

Improving our ability to predict and optimize language recovery after stroke is essential for developing precision, personalized rehabilitation strategies that match patients to the therapies most likely to benefit them ^1–3^. Aphasia severity and recovery potential have long been known to be dependent on stroke lesion size and distribution and more recent work has expanded our understanding of the effects of individual patient factors such as small vessel disease burden, brain age, and structural and functional neuroimaging markers ^4–18^. Together, these findings underscore that post-stroke aphasia (PSA) reflects not only focal damage to the language network but also widespread disruption of structural and functional networks.

A substantial body of neuroimaging research has shown that impairments in PSA arise from both direct lesion effects and dysfunction in remote, structurally connected regions.^9,19–24^ Lesion-induced disconnection has been repeatedly implicated in aphasia severity and recovery trajectories^14,25–27^. Resting-state and task-based functional MRI studies further demonstrate that alterations in functional connectivity, including within left perisylvian language regions as well as distributed frontoparietal and default-mode systems, are associated with both baseline language impairment and recovery potential^9,11,15,21,23,24,28^. These findings highlight the importance of understanding PSA through the lens of large-scale network disruption rather than isolated damage to specific language regions.

Motivated by this network perspective, several groups have used machine-learning models to predict aphasia outcomes from neuroimaging features. Lesion-based and connectome-based approaches have achieved moderate accuracy, and multimodal frameworks that integrate lesion maps, structural connectivity, and functional measures^29,30^ have reported further gains, supporting the idea that different modalities carry complementary information^25,29–32^. More recently, Hu et al.^33^ used a rigorously nested SVR–RFE pipeline to evaluate lesion, structural, and functional connectivity features, and showed that multimodal combinations, particularly those incorporating structural disconnection, functional connectivity, and graph-theoretic measures, yield the strongest predictions of aphasia severity. However, these studies do not address whether the apparent benefits of multimodality reflect truly unique information in each imaging modality, or whether much of the predictive signal is already implicit in the lesion distribution and could be recovered through more principled or mechanistically grounded analyses. Moreover, multimodal MRI acquisition and preprocessing remain resource-intensive, limiting their clinical scalability.

Computational models of large-scale brain dynamics offer a complementary approach for linking structural damage to network-level alterations in neural activity. Neural mass and mean-field models, such as the Wong–Wang framework and subsequent large-scale implementations, approximate population activity within each cortical region and simulate interactions constrained by a structural connectome. ^34–38^ Although these models were originally developed to capture large-scale dynamics in normative, non-lesioned brains, lesion-informed extensions enable simulations tailored to individual stroke patients. Lesion-informed simulations can generate individualized estimates of how structural disruption alters large-scale interactions, capturing changes in activity, inter-regional influence, and deviations from normative connectivity patterns that may not be directly observable from empirical imaging.

To operationalize these ideas, we developed REWIRED, a lesion-informed cortical dynamics framework that incorporates individual lesion information to generate subject-specific estimates of lesion-driven network disruption. We first evaluated whether simulation-derived functional connectivity captured patient-specific variation in empirical functional connectivity beyond lesion burden and structural disconnection alone. We additionally developed a set of lesion-derived, structural, and simulation-derived metrics to predict aphasia severity in chronic PSA. We hypothesized that lesion-informed simulations would capture biologically meaningful variation in functional network organization and that simulation-derived features would improve prediction performance beyond lesion volume, lesion distribution, and structural connectivity alone.

Accordingly, the present study addressed two aims: Aim 1. Evaluate whether lesion-informed cortical dynamics simulations capture patient-specific variation in empirical functional connectivity beyond lesion burden and structural disconnection alone, and Aim 2. Test whether simulation-derived network measures improve prediction of chronic aphasia severity beyond lesion-distribution and structural-connectivity predictors.

## 2. Methods

In this study, we developed REWIRED, a lesion-informed cortical dynamics framework that integrates patient-specific lesion topology with normative structural connectivity to model large-scale network disruption after stroke, and evaluated it using two complementary analyses: (1) validation of simulation-derived functional connectivity against empirical functional connectivity in individuals with chronic post-stroke aphasia, and (2) prediction of chronic aphasia severity, operationalized using the Western Aphasia Battery-Revised Aphasia Quotient (WAB-AQ). An overview of the methods is provided in Figure 1. The major components of the REWIRED pipeline are summarized below with emphasis on the aspects most relevant to the present analyses. Briefly, REWIRED combines lesion-informed structural disconnectome estimation with large-scale cortical dynamics simulations to model the network-level impact of stroke. Its major components include generation of a probabilistic lesion representation, disconnectome estimation, and computational simulation of neural dynamics. Here, we summarize these methods with the goal of providing sufficient detail for evaluating and interpreting the WAB-AQ prediction results.

**Figure 1.**
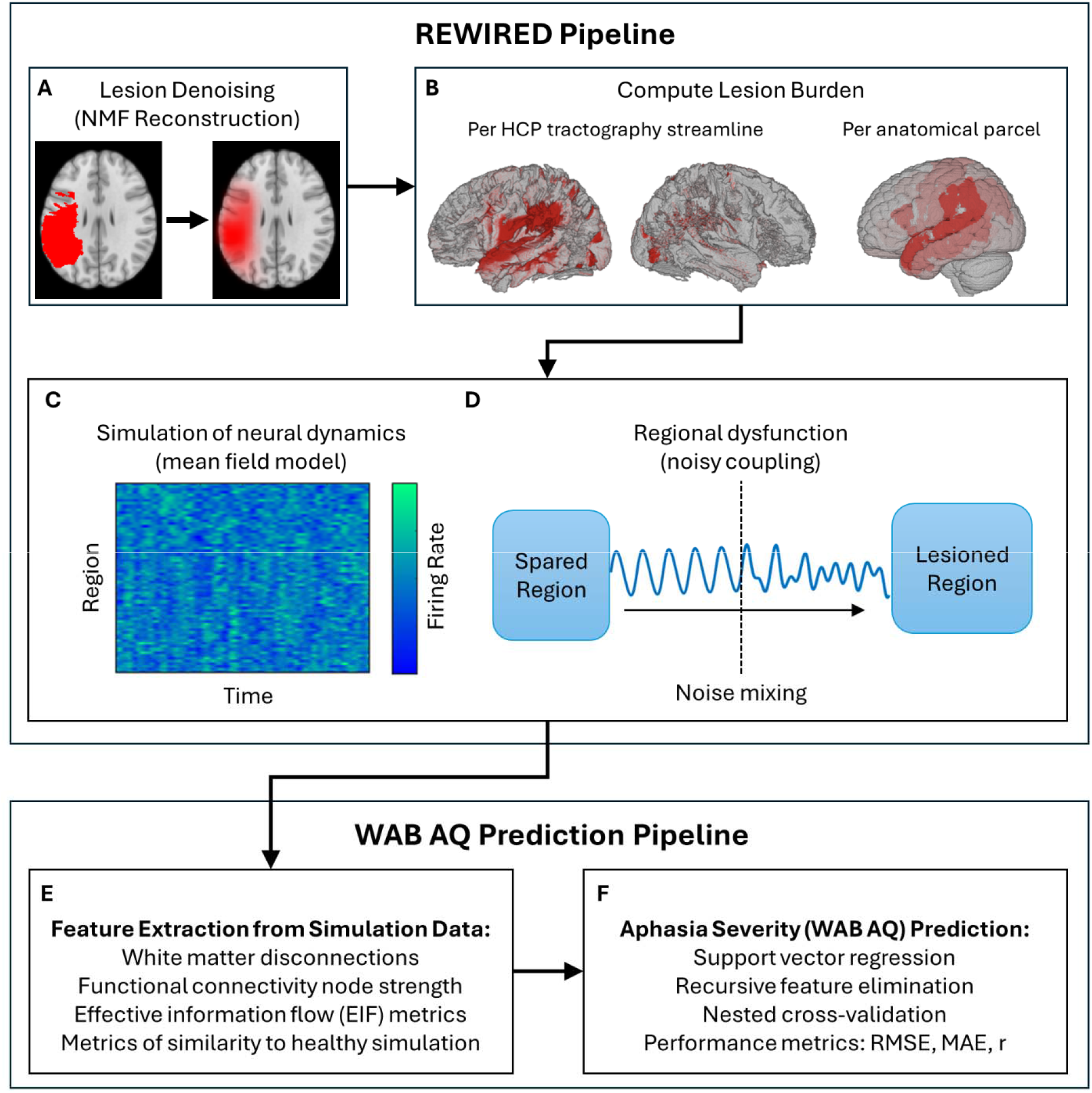
Overview of methods for prediction of aphasia severity using lesion distribution-based personalized simulation of cortical neural activity. **A**. Lesion denoising was achieved by projecting each manual lesion map onto NMF-derived spatial components and reconstructing it as a weighted sum of those components. **B**. Disconnectomes and parcel lesion burden were obtained by overlaying NMF-reconstructed lesion maps on the HCP1065 streamlines and Desikan-Killiany atlas parcels and computing the cumulative proportion lesioned for each streamline or parcel. **C**. large-scale cortical dynamics are simulated using a mean field model, generating firing rate time series for each cortical region. **D**. Parcel lesion burden was used to personalize simulations by modulating inter-regional coupling as a function of regional damage. **E**. Extraction of structural and simulation-derived features, including white matter disconnection-based metrics, functional connectivity node strength, effective information flow measures, and similarity to non-lesioned simulations. **F**. Aphasia severity (WAB-AQ) prediction pipeline. Structural and simulation-derived features are used in a support vector regression model with recursive feature elimination, optimized through nested cross-validation. Prediction accuracy is reported using RMSE, MAE, and Pearson’s r. Abbreviations: NMF = non-negative matrix factorization, HCP = Human Connectome Project, WAB-AQ = Western Aphasia Battery-Revised Aphasia Quotient.

**Figure 2.**
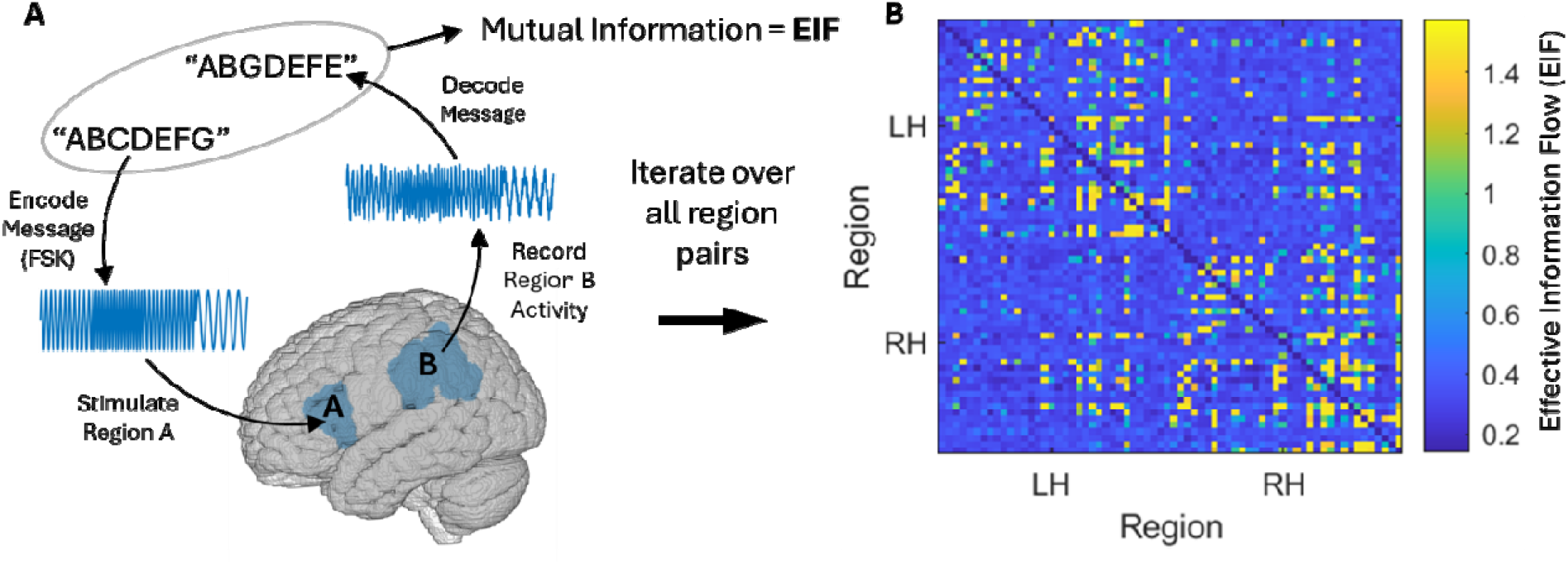
Illustration of effective information flow (EIF), a model-based measure of directed inter-regional communication. **A**. EIF is computed by injecting a temporally varying signal into a single brain region during the neural-dynamics simulation. The resulting activity in all other regions is analyzed to determine how accurately the injected signal can be recovered, yielding a directional EIF value from the transmitting region to each receiving region. **B**. Group-level EIF matrix from the healthy baseline simulation, showing the fidelity of directed information transfer between all pairs of regions in the left (LH) and right (RH) hemispheres. Warmer colors indicate higher decoding accuracy and thus stronger directed information flow.

### 2.1. Participants & Data

#### 2.1.1. Aphasia severity prediction

The primary dataset used in this study came from the Aphasia Recovery Cohort (ARC) dataset which is freely available at https://doi.org/10.18112/openneuro.ds004884.v1.0.1. The dataset consists of deidentified imaging, behavioral measures, and demographic data from a cohort of individuals with chronic post-stroke aphasia collected through multiple studies conducted at the University of South Carolina. For additional details on the dataset, refer to Gibson et al. ^39^.

Participants from the dataset were included if they had a complete set of imaging and behavioral data, defined as availability of T1-weighted and T2-weighted structural MRI, a manually delineated lesion mask, resting-state fMRI (rsfMRI), and a Western Aphasia Battery Aphasia Quotient (WAB-AQ) score. These criteria ensured that all individuals had the full set of inputs required for both the present analyses and for planned extensions of the pipeline that incorporate resting-state functional connectivity. Subjects lacking any of these modalities were excluded. Of the 131 participants in the initial dataset, two participants were excluded due to poor quality and/or failure of MRI preprocessing.

In addition to the ARC dataset, the study made use of group averaged tractography data from the Human Connectome Project (HCP; Van Essen et al., 2012) as well as an HCP-derived structural connectome and optimized simulation parameters from Kong et al. (2021). These data are freely available at https://brain.labsolver.org/hcp_trk_atlas.html and https://github.com/ThomasYeoLab/CBIG/tree/master/stable_projects/fMRI_dynamics/Kong2021_pMFM, respectively. The data were used for baseline healthy comparisons and as a basis from which to generate personalized lesion-modified neural mass models. These methods are described in their respective sections below.

#### 2.1.2. Empirical functional connectivity validation

Resting-state fMRI data used for empirical functional connectivity analyses were obtained from a separate cohort of 25 individuals with chronic post-stroke aphasia collected as part of the Center for the Neurobiology of Language Recovery (CNLR) project^11,41,42^. Images were acquired on Siemens 3T MRI scanners. High-resolution T1-weighted structural images were acquired for anatomical preprocessing and lesion normalization. Functional images were acquired using a gradient-echo T2*-weighted sequence (TR = 2.4 s, TE = 20 ms, flip angle = 90°, voxel size = 1.72 × 1.72 × 3 mm^3^). Structural preprocessing included enantiomorphic replacement of lesioned tissue using the SPM Clinical Toolbox^43^ prior to normalization. Structural and functional preprocessing was then performed using fMRIPrep version 1.3.175 and included motion correction, normalization to MNI152NLin2009cAsym space, tissue segmentation, and registration of functional images to structural anatomy. Lesion voxels were excluded from functional data following normalization. Preprocessed resting-state data were imported into the CONN toolbox^44^ for denoising and functional connectivity estimation. Denoising included motion outlier detection using the Artifact Detection Tool (ART), nuisance regression of motion parameters and white matter/cerebrospinal fluid signals, linear detrending, and temporal band-pass filtering (0.008–0.09 Hz). Mean regional BOLD time series were extracted using the Desikan–Killiany atlas^45^, and empirical functional connectivity matrices were computed as pairwise Pearson correlations between regional time series.

### 2.2. Lesion Mask Normalization

The ARC dataset includes manually segmented lesion masks drawn on each participant’s T2-weighted structural scan. To normalize these masks to MNI space, T2 images were first rigidly coregistered to each participant’s T1-weighted image using the Coregister module in SPM12 ^46,47^, and lesion masks were resliced into T1 space using nearest-neighbor interpolation to preserve binary boundaries. Each T1 image was then normalized to the SPM12 T1 template using the Old Normalize procedure (spm_normalise), which computes affine and nonlinear deformations ^48^. The resulting deformation parameters were applied to the T1-aligned lesion mask using spm_write_sn (nearest-neighbor interpolation), producing a lesion mask in MNI152 space.

### 2.3. REWIRED Pipeline

As shown in Figure 1, this pipeline first involves lesion denoising using Non-Negative Matrix Factorization (NMF) reconstruction. Next, the lesion burden is computed by quantifying the damage relative to Human Connectome Project (HCP) tractography streamlines and anatomical parcels. This lesion information is incorporated into large-scale cortical dynamics simulations to model lesion-dependent disruption of neural activity and inter-regional network interactions. From these simulations, features such as white matter disconnections, functional connectivity node strength, and effective information flow (EIF) are extracted. Finally, these features are used to predict Aphasia Severity (WAB-AQ) using Support Vector Regression (SVR) combined with Recursive Feature Elimination (RFE), validated through nested cross-validation, and evaluated using RMSE, MAE, and correlation coefficients.

#### 2.3.1. Lesion denoising using nonnegative matrix factorization (NMF)

As an alternative to traditional lesion symptomatology approaches, we employed a denoising technique using non-negative matrix factorization (NMF) to reconstruct each subject’s lesion map from a set of latent spatial components that capture the dominant patterns of lesion distribution across the dataset. NMF overcomes traditional statistical limitations of analyzing millions of brain voxels by decomposing complex lesion maps into a small number of biologically interpretable, additive components, such as specific vascular territories, allowing researchers to capture overlapping damage patterns and predict behavioral deficits. We adapted an approach described in Kropp et al. (2025) to apply NMF to region-wise lesion burden data and identify latent stroke patterns shared across this cohort. First, all binary lesion masks were registered to MNI space and combined to define a left-hemisphere analysis mask (since all strokes were left-dominant). To avoid parcellation bias from predefined atlas ROIs, we tiled the brain with uniform, non-overlapping 9-voxel cubes. Cubes were retained if ≥80% of their voxels lay within this left-hemisphere analysis mask. We explored cube sizes and overlap thresholds to maximize coverage of lesioned voxels while keeping the cube count minimal. For each retained cube, we computed participant-wise lesion load and applied NMF to this data to identify 16 spatial lesion patterns (with k=16 picked via the method in Kropp et al., 2025) and corresponding subject weights (how well each pattern described individual participant lesions). To generate denoised lesion maps, each participant’s lesion was reconstructed as the dot product of the spatial lesion patterns and that participant’s NMF scores.

The resulting continuous maps were projected back to MNI space to yield probabilistic lesion maps, where higher values indicate greater confidence that the voxel is truly lesioned. These reconstructions reduced inconsistencies in manual segmentations while preserving the dominant lesion topology captured by the latent spatial stroke patterns.

#### 2.3.2. Regional lesion burden and structural disconnectomes

Neural dynamics simulations (described in MFM section below) used personalized structural disconnectomes and regional lesion burden computed from each subject’s lesion data. Parcel-level lesion burden was obtained by overlaying each subject’s NMF-reconstructed lesion map onto the Desikan–Killiany (DK) anatomical parcellation ^45^. For each parcel, lesion burden was calculated as the sum of lesion-map voxel intensities divided by the total number of voxels in the parcel.

To generate structural disconnectomes, we first constructed a baseline structural connectome using the DK atlas and the HCP1065 population-averaged tractography atlas ^49^. Using a custom MATLAB implementation, the DK atlas was overlaid on the HCP1065 tractogram, and each streamline’s originating and terminating parcels were identified to assemble the baseline structural connectivity matrix using streamline counts. Next, the HCP1065 atlas was overlaid with each subject’s NMF-reconstructed lesion map to compute cumulative lesion burden for each streamline. Proportion spared for each streamline was defined as 1−lesion burden. The lesioned structural connectome was then generated by repeating the streamline-parcel assignment but weighting each streamline’s contribution to the connection strength by its proportion spared. Finally, each subject’s structural disconnectome was computed as (baseline structural connectivity – lesioned structural connectivity) / baseline structural connectivity.

#### 2.3.3. Mean field model (MFM) with lesion effects

Cortical neural dynamics were simulated for both the normative non-lesioned and subject-specific lesioned conditions using the mean field model (MFM) described in Wang et al. ^37^ and updated by Kong et al. ^38^ Briefly, the MFM is a large-scale, rate-based dynamical model in which each cortical region’s activity is determined by its (1) recurrent input, (2) inter-regional input, and (3) external input:

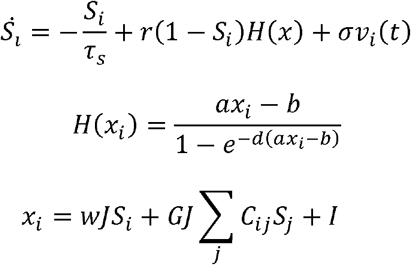

Here, 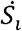 is the synaptic gating variable, *H*(*x*_*i*_) is the population firing rate function, and *x*_*i*_ is the total input current to region *i*. The total input current depends on the recurrent connection strength *w*, synaptic coupling constant *J*, the global scaling factor *G*, and the structural connectivity weights *C*_*ij*_, as well as the external input *I*. Parameters *a, b*, and *d* define the sigmoidal input–output transfer function *H*(*x*_*i*_). The parameters *r* and *τ*_*s*_ govern the kinetics of synaptic activity, and *σv*_*i*_ (*t*) represents Gaussian noise with amplitude σ. Kong et al.’s (2021) extension incorporates region-specific synaptic parameters, obtained by parameterizing the first principal component of resting-state functional connectivity in HCP data. The present study uses the regional and global parameter values reported in Kong et al. ^38^, reproduced in Supplementary Table 1.

**Table 1.** Linear mixed-effects model predicting edgewise empirical functional connectivity from simulation-derived functional connectivity, edgewise structural disconnection, and regional lesion burden. Random intercepts were included for participant and edge identity to account for repeated observations across subjects and network edges. Simulation-derived functional connectivity significantly predicted empirical functional connectivity after accounting for lesion burden and structural disconnection, supporting the biological validity of the REWIRED framework.

| Predictor | Beta | SE | t | p | 95%_CI |
| --- | --- | --- | --- | --- | --- |
| Intercept | 0.0232 | 0.0062 | 3.75 | 1.80E-04 | [0.0110,0.0353] |
| Simulated FC | 0.1365 | 0.0218 | 6.27 | 3.74E-10 | [0.0938,0.1792] |
| Lesion burden node 1 | -0.0148 | 0.0065 | -2.28 | 0.0226 | [-0.0275,-0.0021] |
| Lesion burden node 2 | 0.0245 | 0.0092 | 2.65 | 0.008 | [0.0064,0.0426] |
| Edgewise disconnection | -0.0119 | 0.007 | -1.72 | 0.0861 | [-0.0256,0.0017] |
| **Random intercept SD:** |  |  |  |  |  |
| Subject ID | 0.019 |  |  |  |  |
| Edge ID | 0.1774 |  |  |  |  |
| Residual error | 0.1926 |  |  |  |  |

Abbreviations: FC = functional connectivity; SE = standard error; CI = confidence interval; SD = standard deviation.

The REWIRED framework extends the baseline cortical dynamics model by incorporating lesion-dependent disruption of inter-regional communication. Regional lesion burden modulates the influence of structurally mediated inputs, allowing simulations to reflect progressive degradation of large-scale network interactions following stroke.

### 2.4. Validation of simulation-derived functional connectivity

To evaluate whether REWIRED simulations captured patient-specific variation in empirical functional connectivity, we compared simulation-derived and empirical functional connectivity matrices in the CNLR cohort. For each participant, empirical functional connectivity was represented as a 68 × 68 Desikan–Killiany parcel-wise matrix, and simulation-derived functional connectivity was extracted from the corresponding lesion-informed cortical dynamics simulation. Edgewise data were vectorized using the upper triangle of each matrix. We then fit a linear mixed-effects model predicting empirical functional connectivity from simulation-derived functional connectivity while adjusting for edgewise structural disconnection and lesion burden in the two regions defining each edge. Random intercepts for participant and edge identity were included to account for repeated observations across participants and edges.

### 2.5. Feature Extraction and Aphasia Severity Prediction

Using the disconnectome and simulation data generated as described above, a set of features characterizing neural dynamics were extracted. These features, described in greater detail below, were combined with lesion distribution-based features in a machine learning pipeline using support vector regression (SVR) and recursive feature elimination (RFE) inside nested cross-validation.

#### 2.5.1. Features

The complete feature set included three feature classes: lesion distribution-based features, structural disconnectome-based features, and simulated neural dynamics-based features. Lesion distribution features included lesion volume and the 16 NMF-derived lesion patterns (“atoms”). Disconnectome features included regional structural connectivity strength computed on the lesion-modified HCP structural connectome, as well as network- and region-level “similarity” metrics as described below. Neural dynamics features consisted of global and regional functional connectivity-based metrics and effective information flow metrics. A complete list of features is provided in Supplementary Table 2. Effective information flow, a novel metric developed as part of this work, and “similarity” metrics, which measure the degree to which a subject’s lesion-modified connectome or neural dynamics deviate from the non-lesioned condition, are described in the following sections.

**Table 2.**
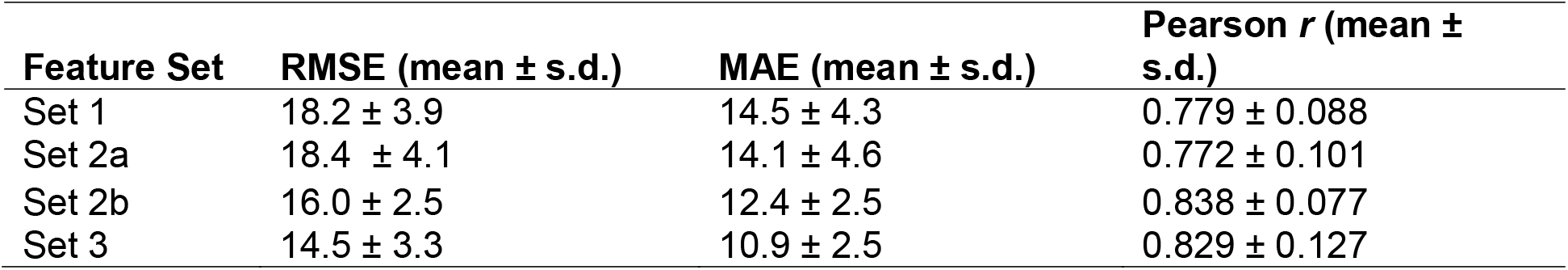
Nested cross-validated prediction performance for each feature set.

| Feature Set | RMSE (mean ± s.d.) | MAE (mean ± s.d.) | Pearson $r$ (mean ± s.d.) |
| --- | --- | --- | --- |
| Set 1 | 18.2 ± 3.9 | 14.5 ± 4.3 | 0.779 ± 0.088 |
| Set 2a | 18.4 ± 4.1 | 14.1 ± 4.6 | 0.772 ± 0.101 |
| Set 2b | 16.0 ± 2.5 | 12.4 ± 2.5 | 0.838 ± 0.077 |
| Set 3 | 14.5 ± 3.3 | 10.9 ± 2.5 | 0.829 ± 0.127 |

##### 2.5.1.1. Effective information flow (EIF)

To quantify directed inter-regional communication, we developed effective information flow (EIF), a simulation-derived measure based on the propagation and recovery of temporally varying signals through the modeled cortical network. EIF quantifies how faithfully information introduced into one region can be recovered from activity in another region, yielding a directed estimate of inter-regional communication fidelity.

Directional EIF for a given region pair was defined as the mutual information between the original message and the decoded message, an information-theoretic metric that measures the dependence between the two signals and quantifies the fidelity with which information injected at the source is recovered at the target. Repeating this procedure with each region as the message source yielded a directed 68×68 EIF matrix summarizing the pattern of information flow throughout the cortical network.

Several summary features are derived from this matrix. EIF_rx_ strength for a node is computed as the sum of its incoming EIF values, representing the extent to which messages injected into other regions can be decoded from that node’s activity. Conversely, EIF_tx_ strength is the sum of a node’s outgoing EIF values, reflecting the extent to which messages injected into that node can be decoded from the activity of all other regions. EIF local efficiency quantifies the efficiency of directed communication among a node’s immediate neighbors. At the global level, EIF_RX_ strength similarity and EIF_TX_ strength similarity measure how closely a lesioned simulation preserves normal patterns of directed communication. Higher similarity indicates more preserved directed communication structure despite the lesion. The following section describes how these and other global and regional “similarity” metrics were computed.

##### 2.5.1.2. Metrics of similarity to baseline healthy condition (“similarity metrics”)

In addition to the metrics discussed above, we also calculated several similarity metrics to quantify how closely each lesioned simulation resembled the healthy baseline simulation. Both global and node-level similarity metrics were computed. To obtain baseline values, the MFM was run six times, and the mean of the resulting edge-level metrics was used as the baseline reference.

For global metrics, a single value per node (e.g., FC strength) was computed for both the baseline and lesioned simulations, and similarity was defined as the Pearson correlation between the two sets of values. For node-level metrics, a vector of values per node was extracted (e.g., each node’s structural connectivity weights to all other nodes), and similarity was again computed as the Pearson correlation between the baseline and lesioned simulations.

##### 2.5.1.3. Correlational analysis of node-level features

To characterize relationships among the node-level features described above, we computed pairwise Pearson correlations between the group-averaged values of each metric across the 68 cortical regions. Metrics included lesion burden (proportion lesioned per parcel) and structural connectivity, functional connectivity, EIF_RX_ and EIF_TX_ strength. This analysis provided a quantitative assessment of the extent to which each feature reflected the spatial distribution of focal damage and the degree to which different dynamic and structural measures captured overlapping or distinct aspects of lesion-induced network disruption.

#### 2.5.2. Prediction of WAB-AQ

Prediction of WAB-AQ was performed using a supervised machine learning framework adapted from Hu et al.^33^ To evaluate the relative contribution of lesion-, disconnection-, and simulation-derived information, we compared four nested feature sets: Set 1 (lesion volume + NMF loading scores), Set 2a (Set 1 + disconnectome-based metrics), Set 2b (Set 1 + simulation-derived metrics), and Set 3 (Set 1 + disconnectome-based metrics + simulation-derived metrics). To limit the size of feature sets, only left hemisphere region-level features were included as right hemisphere regions were expected to show less intersubject variation and carry less predictive signal. Global features were computed on the full bilateral set of regions and thus capture large scale network patterns including right hemisphere information.

All models were trained and evaluated using a nested cross-validation (CV) procedure designed to ensure unbiased performance estimates and strict separation of feature selection from test data. Support Vector Regression (SVR) models with both linear and radial basis function (RBF) kernels were examined. The full dataset was partitioned into 11 outer folds, each serving once as a held-out test set while the remaining data formed an outer training set. To conduct feature selection and hyperparameter tuning without contaminating test data, each outer training set was further partitioned into 10 inner folds. All feature selection and model optimization steps were carried out exclusively within these inner folds, and the held-out outer fold was used only for final performance evaluation.

Feature selection was performed using recursive feature elimination (RFE), consistent with Hu et al.^33^ For each inner training fold, a linear SVR model was fit to compute coefficient-based feature importance scores (all features were standardized before model fitting). At each RFE iteration, the bottom 40% of features by importance were removed and the model was refit until a prespecified number of features, k, remained. This procedure was repeated across the 10 inner folds of the nested cross-validation, producing a count of how often each feature was retained among the top k. For each candidate k in the range 5–18, the k features with the highest selection frequencies across the 10 inner folds were selected as the feature set for that outer fold’s final model, which was then used to predict the held-out test data.

For each feature-set size k, both SVR models (linear and RBF) were trained and tuned using the 100 inner validation–training splits, with hyperparameters selected based on average inner-fold RMSE. The value of k and model configuration yielding the lowest mean validation RMSE were carried forward to the outer loop. The selected model was then retrained on the entire outer training fold and evaluated on the held-out outer test fold. Final performance was summarized as the average test-set RMSE, MAE, and cross-validated R^2^ across all 11 outer folds.

All models were implemented in Python using scikit-learn (Pedregosa et al., 2011). Methodological details on the nested CV framework, RFE procedure, and SVR optimization are described in Hu et al.^33^

## 3. Results

Using the REWIRED framework described above, we derived lesion-, disconnectome-, and simulation-derived features characterizing structural connectivity, functional connectivity, and effective information flow. We first evaluated whether simulation-derived functional connectivity captured biologically meaningful variation in empirical functional connectivity beyond lesion burden and structural disconnection alone (Aim 1). We then evaluated whether these simulation-derived network measures improved prediction of chronic aphasia severity (WAB-AQ) beyond lesion- and disconnectome-derived features (Aim 2).

### 3.1. Validation of simulation-derived functional connectivity (Aim 1)

To evaluate whether lesion-informed cortical dynamics simulations captured biologically meaningful variation in empirical functional connectivity, we fit a linear mixed-effects model predicting edgewise empirical functional connectivity from simulation-derived functional connectivity while adjusting for edgewise structural disconnection and lesion burden in the two regions defining each edge. Random intercepts for participant and edge identity were included to account for repeated observations across participants and edges.

Simulation-derived functional connectivity significantly predicted empirical functional connectivity after accounting for regional lesion burden and edgewise structural disconnection (β = 0.137, t = 6.27, p = 3.74 × 10^− 1^□). In contrast, edgewise structural disconnection was not a significant predictor after accounting for simulation-derived functional connectivity (p = 0.086). These findings indicate that the lesion-informed simulations captured patient-specific distributed patterns of functional network disruption beyond the direct effects of focal tissue damage and structural disconnection alone. Together, these results support the biological validity of the REWIRED framework and suggest that the simulations capture patient-specific alterations in large-scale functional network organization.

### 3.2. Characterizing simulation-derived network disruption metrics

We developed a set of nine global, three network-level, and six regional simulation-derived metrics (68 features per regional metric) that characterize lesion-driven alterations in structural connectivity, functional connectivity, and effective information flow, along with additional metrics that quantify the degree to which each participant’s simulated dynamics diverged from the non-lesioned normative simulation. To qualitatively assess whether these measures provide independently informative and neuroanatomically interpretable markers of network disruption, we examined the spatial distribution of the group-level mean for each regional metric (Figure 3). As expected, regional lesion burden and structural connectivity strength reductions were largely confined to left perisylvian and perirolandic regions. In contrast, functional connectivity and effective information flow metrics, which depend on simulated neural dynamics rather than direct structural damage, showed more spatially distributed abnormalities, with deviations from the non-lesioned simulation extending into regions of the right hemisphere. Notably, aside from a modest left-lateralized asymmetry, the spatial distributions of EIF_TX_ and EIF_RX_ did not align with the lesion-burden map: regions showing abnormalities in directed information flow were often not those directly damaged by the stroke, indicating that these metrics are capturing large-scale, network-level consequences of focal injury, consistent with empirical findings in the functional connectivity literature.^50–52^ Finally, the distinct spatial patterns of EIFTX and EIFRX suggest that they are differentially affected by lesion-induced disconnection and regional dysfunction, and may capture complementary dimensions of network impairment. For example, several central lobe regions show increased EIF_RX_ but decreased EIF_TX_, indicating that these regions become stronger receivers but weaker transmitters of directed information flow, an asymmetry that is evident in Figure 3.

**Figure 3.**
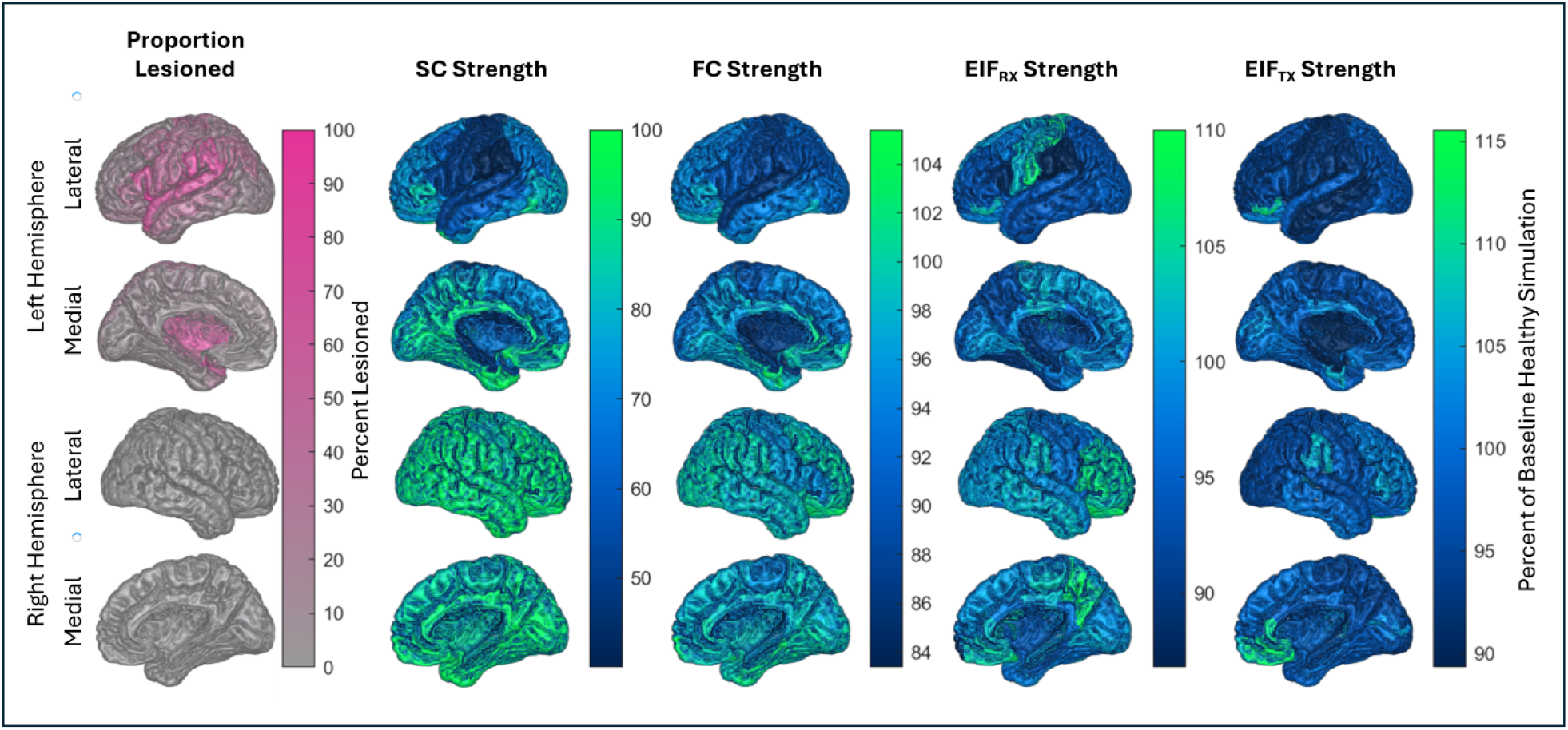
Spatial distribution of regional lesion burden, structural and functional connectivity strength, and effective information flow (EIF) metrics. Group-level mean maps are shown for regional proportional lesioned, structural connectivity strength, functional connectivity strength, and incoming (EIF_RX_) and outgoing (EIF_TX_) effective information flow, expressed relative to the non-lesioned baseline simulation. Lesion burden and reductions in SC strength were most pronounced in left perisylvian and perirolandic regions. In contrast, functional connectivity strength and EIF measures displayed more spatially distributed abnormalities. Notably, the spatial patterns of EIF_RX_ and EIF_TX_ were less directly related to lesion distribution indicating large-scale network consequences of focal injury. The distinct distributions of EIF_RX_ and EIF_TX_ suggest asymmetries between information-receiving and information-transmitting capacities after stroke.

Correlation analyses (Figure 4) showed that lesion burden was highly correlated with structural and functional connectivity strength. Consistent with their limited spatial resemblance to the lesion distribution (Figure 3), both EIF_RX_ and EIF_TX_ strength were only moderately correlated with lesion burden and with structural and functional connectivity. This pattern suggests that EIF metrics capture large-scale lesion-induced network dysfunction that is not simply a reflection of focal damage. Moreover, EIF_RX_ and EIF_TX_ strength were only weakly correlated with each other. EIF_RX_ indexes the extent to which information arriving from the rest of the brain is expressed in a region’s dynamics, capturing its susceptibility to upstream disruptions, whereas EIF_TX_ reflects how strongly perturbations originating in that region influence the broader network, capturing its downstream broadcast capacity. Because these direction-specific properties depend on partially distinct pathways and dynamical interactions, it is expected that they correlate only weakly, indicating that they capture complementary aspects of network organization. Taken together, these results demonstrate that simulation-derived metrics quantify distinct and complementary dimensions of lesion-induced network disruption that are not explained by focal structural damage alone.

**Figure 4.**
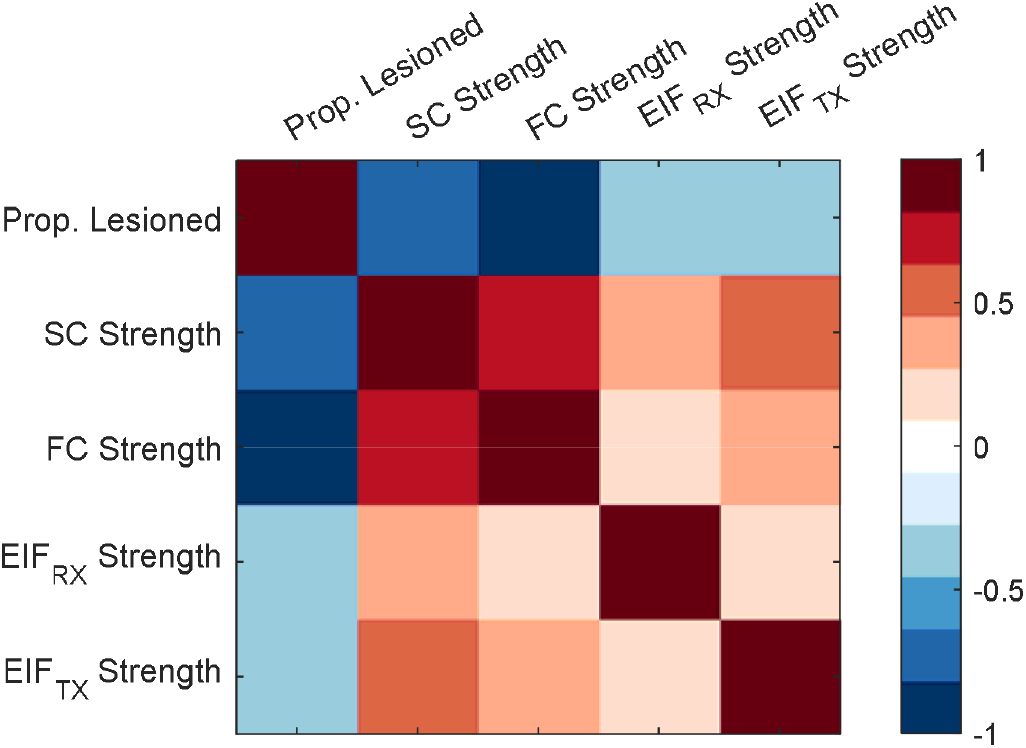
Correlation matrix among lesion burden, connectivity strengths, and effective information flow (EIF) measures. Lesion burden was strongly correlated with structural and functional connectivity strength, whereas EIF_RX_ and EIF_TX_ strengths showed only moderate correlations with these measures and with lesion burden. EIF_RX_ and EIF_TX_ were weakly correlated with each other, highlighting their complementary roles in capturing directed aspects of network disruption after stroke.

### 3.3. Aphasia severity prediction (Aim 2)

To evaluate the respective contributions of each of the three feature classes to aphasia severity prediction, we compared four nested feature sets: Set 1 (lesion volume + NMF loading scores), Set 2a (Set 1 + disconnectome-based metrics), Set 2b (Set 1 + simulation-derived metrics), and Set 3 (Set 1 + both disconnectome- and simulation-derived metrics). Prediction performance was relatively poor for Sets 1 and 2a, with a marked improvement when simulation-derived features were added in Sets 2b and 3. Set 3 achieved the lowest overall prediction error (RMSE = 14.5, MAE = 10.9, r = 0.83), indicating that simulation-derived metrics provide predictive value beyond lesion characteristics and structural connectivity features. Performance metrics for each set are shown in Table 1, and Supplementary Table 3 lists the selected features for each set.

Feature-selection stability differed markedly across the feature sets and closely mirrored their cross-fold predictive performance. Sets 1 and 2a showed relatively unstable feature selection: among the features selected in at least one outer fold, 33% (Set 1) and 42% (Set 2a) were selected in fewer than 20% of folds. This instability was accompanied by poor predictive performance across the 11 outer folds. In contrast, Set 3 exhibited both high stability and strong cross-fold performance. The top 16 features in Set 3 were selected in more than 90% of folds, and only 6% of features selected at least once appeared in fewer than 20% of folds. Consistent with this stability, Set 3 achieved the best performance in 9 of the 11 outer folds. Density plots illustrating feature selection stability differences are shown in Figure 5.

**Figure 5.**
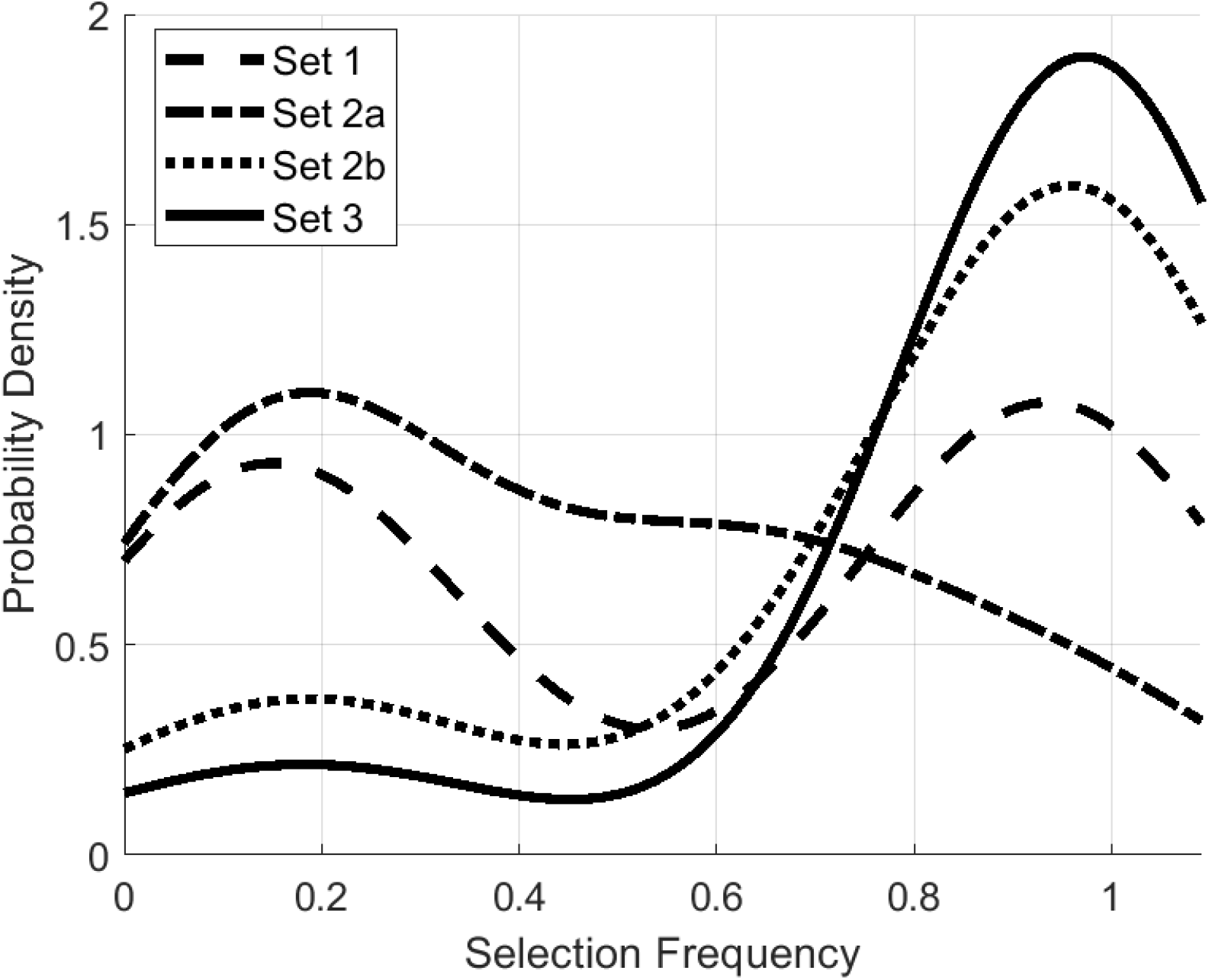
Density plots showing a comparison of feature selection stability. Greater density near 1 (Sets 2b and 3) indicates high stability with many features selected in all or nearly all folds. High density below 0.4 (Sets 1 and 2a) indicates feature selection instability with many features selected in fewer than 40% of folds. Only features that were selected in at least one fold were included in the stability analysis.

## 4. Discussion

In this study, we developed and evaluated REWIRED, a lesion-informed cortical dynamics framework designed to model large-scale network disruption after stroke and generate simulation-derived measures of functional brain organization. Using REWIRED, we demonstrated that simulation-derived functional connectivity captured biologically meaningful variation in empirical functional connectivity and that simulation-derived network measures substantially improved prediction of chronic aphasia severity beyond lesion- and disconnectome-derived features alone. Across analyses, each successively added feature class improved prediction performance, with the full feature set outperforming models based solely on lesion anatomy and achieving accuracy competitive with current multimodal imaging methods.

Below, we first situate the predictive performance of our approach in the context of prior work, and then discuss each feature class in turn, interpreting its relevance for understanding network disruption after stroke and improving individualized outcome prediction. Because the spatial distributions and correlation patterns of these metrics (Aim 1) directly inform their predictive utility and neurobiological interpretation, we integrate discussion of Aim 1 and Aim 2 findings, situating the observed patterns within frameworks of diaschisis and large-scale network disruption.

### 4.1. Validation of simulation-derived functional connectivity

A central finding of the present study is that simulation-derived functional connectivity significantly predicted empirical functional connectivity even after accounting for lesion burden and structural disconnection. This suggests that the REWIRED simulations capture patient-specific distributed patterns of functional network organization beyond the direct effects of focal tissue damage alone. Importantly, the empirical functional connectivity analyses were performed using the CNLR cohort which was independent of the aphasia severity prediction dataset, providing converging support for the biological validity of the framework. Together, these findings indicate that lesion-informed cortical dynamics simulations can recover meaningful aspects of large-scale functional network disruption from structural lesion information alone.

### 4.2. Predictive performance

Prior work using machine-learning models to predict aphasia severity from neuroimaging has shown moderate accuracy. A broad body of work, including lesion-load models,^53–56^ structural and functional connectome–based approaches,^30^ and multimodal stacking frameworks,^29^ has demonstrated that multimodal data can incrementally improve prediction, though gains have often been modest^25,30,32^. Among the most comparable recent studies, Kristinsson et al.^30^ reported r = 0.67 for WAB-AQ using a multimodal SVR model integrating lesion load, diffusion metrics, cerebral blood flow, and task-based fMRI, while Hu et al.^33^ achieved RMSE = 15.82 and r = 0.73 using a rigorously nested SVR–RFE framework combining lesion characteristics, structural connectivity, and functional connectivity. In the present study, the best REWIRED-derived feature set surpassed these benchmarks (RMSE = 14.5; r = 0.83), despite relying solely on lesion-distribution inputs with all functional connectivity and directed-communication features generated by the REWIRED simulation rather than acquired through multimodal MRI. The strong feature-selection stability observed for the highest-performing models further argues against substantial overfitting despite the high dimensionality of the feature space. These findings suggest that lesion-informed neural-dynamics modeling can recover much of the behaviorally relevant network information typically obtained through multimodal imaging, while relying primarily on routinely acquired structural imaging and therefore potentially offering a more clinically scalable approach to individualized outcome prediction. Importantly, this approach does not preclude multimodal augmentation; future work combining REWIRED simulation-derived metrics such as effective information flow and simulated functional connectivity with diffusion and functional imaging may yield additional gains in predictive accuracy.

### 4.3. Feature class contributions

Each feature class contributes a distinct layer of network information: lesion-distribution features capture the focal anatomical injury, disconnectome metrics describe its structural consequences on white-matter pathways, and simulation-derived measures characterize the resulting alterations in large-scale neural dynamics. Together, these complementary representations allow us to evaluate how different components of the post-stroke network contribute to aphasia severity.

#### 4.3.1. Lesion distribution-based features

Lesion distribution–based features (Set 1) included lesion volume and 16 NMF loading scores that quantify the contributions of specific spatial patterns to each participant’s lesion. Prior work using NMF^18^ has shown that such data-driven lesion patterns capture clinically meaningful anatomical variation and are significantly associated with aphasia severity, even after adjusting for lesion volume. Consistent with these findings, several NMF-derived lesion patterns (“atoms”) were consistently selected by our machine-learning models even when more comprehensive feature sets were considered, indicating that they carry strong predictive signal. In particular, atoms 4, 5, and 12 were selected from Set 3 in all 11 folds. Supplementary Figure 1 visualizes these spatial patterns overlaid on an MNI T1 template.

Atom 4 predominantly involves the left inferior frontal gyrus (pars opercularis/triangularis), with extension into the middle frontal gyrus, adjacent sulcal regions, and underlying frontal white matter. This pattern closely aligns with the MCA superior-division territory. Atom 5 primarily affects left subcortical white matter, including the corona radiata, centrum semiovale, and internal capsule, with involvement of long-range frontoparietal association pathways; its distribution is most consistent with the MCA inferior division. Atom 12 involves temporoparietal deep white matter structures, including posterior corona radiata, centrum semiovale, the arcuate fasciculus, and superior longitudinal fasciculus, with additional involvement of parietal interhemispheric callosal fibers. This pattern corresponds to deep MCA territory and is a classic location for hypertensive hemorrhagic strokes. Taken together, these atoms provide reasonable estimates of the structural regions most critically involved in language processing and recovery, reflecting major perisylvian cortical areas and the white-matter pathways that support them.

Overall, these NMF-derived lesion patterns are consistent with lesion distributions commonly associated with aphasia. Atoms 4 and 12 align closely with classic nonfluent/Broca’s and fluent/Wernicke’s aphasia syndromes, respectively, while atom 5 corresponds to a subcortical nonfluent aphasia. These patterns highlight core structural regions capable of supporting language recovery, insofar as damage to these hubs and their connecting pathways disrupts both perisylvian language circuitry and broader cognitive systems that scaffold communication. Although these primary lesion-distribution features are not sufficient for high-accuracy prediction of aphasia severity, they capture much of the core predictive signal, which is further refined and augmented by the disconnectome-based and simulation-derived metrics discussed below.

#### 4.3.2. Structural disconnectome-based features

Feature Set 2a augmented lesion-distribution measures with structural disconnectome metrics derived from probabilistic NMF-reconstructed lesion maps. Disconnectome features alone (Set 2a) showed poor predictive performance, suggesting limited standalone value. Nevertheless, their contribution became evident when considered alongside simulation-derived metrics: Set 3, which integrates disconnectome metrics with simulation-derived and lesion-distribution features, substantially outperformed Set 2b (lesion + simulation features). The full model reduced RMSE from 16.0 to 14.5, achieved lower error in 10 of 11 outer folds, and yielded a mean per-fold RMSE improvement of 1.46. Feature selection stability also improved modestly (Figure 5). These results suggest that the predictive contribution of structural disconnection becomes fully expressed only when embedded within a dynamical modeling framework that captures the functional consequences of these structural perturbations. In this context, disconnectome features likely reflect large-scale network degradation extending beyond perisylvian language pathways to include domain-general systems involved in attention, cognitive control, and sensorimotor integration.

Consistent with this interpretation, several disconnectome-based features were consistently selected in the final comprehensive model (Set 3), including measures centered in left supramarginal gyrus (SMG), postcentral gyrus (PCG), and rostral anterior cingulate cortex (rACC). Notably, only one of these regions (SMG) is part of the classical perisylvian language network, underscoring that predictive structural disconnection effects extend beyond focal language cortex to encompass broader networks supporting cognitive and sensorimotor processes relevant for speech and language performance.

Overall, these findings indicate that structural disconnection contributes complementary predictive value that emerges most clearly when combined with neural-dynamics simulations. The integration of these modalities in Set 3 demonstrates that both focal and distributed forms of network injury are important for explaining variability in aphasia severity, and that neither structural nor dynamical features alone provide a complete account of post-stroke language impairment.

#### 4.3.3. Simulation-derived features

Simulation-derived features (Set 2b) markedly improved prediction accuracy relative to lesion-distribution and disconnectome features alone, reducing RMSE from 18.2–18.4 in Sets 1 and 2a to 16.0 and increasing mean r to 0.838. These gains demonstrate that personalized neural-dynamics simulations capture aspects of network function that are not represented by static structural or lesion-derived metrics. Importantly, these improvements were achieved despite using only subject-specific binary lesion maps as input, with all functional and dynamical features generated by the simulation rather than acquired through multimodal MRI. This underscores that the parameterized mean field model embedded within the REWIRED framework is able to translate focal structural damage into biologically plausible large-scale functional disruptions. This supports the broader conclusion that lesion-induced alterations in cortical dynamics carry substantial explanatory power for behavioral impairment.

The contribution of simulation-derived measures became even more evident in Set 3, where combining dynamical features with disconnectome metrics yielded the best overall performance (RMSE = 14.5; MAE = 10.9). This improvement over Set 2b (RMSE = 16.0) indicates that structural and functional information are complementary, with simulations providing a functional readout of how disconnection alters interregional coupling and network-level communication. This finding aligns with recent computational and neuroimaging work showing that the temporal properties of large-scale network dynamics, such as criticality, flexible network reconfiguration, and information-processing capacity, carry behaviorally relevant information beyond what static structural or functional measures can capture.^57–59^ The selected simulation features in both Sets 2b and 3 included a mix of core language-related regions (e.g., inferior frontal gyrus, inferior temporal gyrus) and domain-general areas implicated in attention, cognitive control, and integrative processing (e.g., cingulate, orbitofrontal cortex, cuneus, frontopolar cortex). This distributed pattern is consistent with the well-established view that aphasia severity reflects not only damage to classic perisylvian language cortex but also disturbances in the broader networks that support goal-directed behavior, working memory, and sensorimotor integration.^20,56,60^ The substantial contribution of these domain-general regions, despite many not being directly lesioned, highlights the strength of a dynamical approach for capturing remote effects and chronic diaschisis that are difficult to infer from structural data alone.

Perhaps most notable is that simulation-derived measures repeatedly outperformed both lesion-based and disconnectome-derived features when considered independently, despite requiring no additional patient-specific neuroimaging. The ability of REWIRED simulations to generate meaningful functional signatures from simple lesion masks supports the validity of the underlying parameterized mean field model and provides strong converging evidence that it captures neurobiological properties relevant to language processing and language recovery. These findings reinforce the conclusion that personalized neural-dynamics modeling represents a powerful and clinically scalable approach for linking structural injury to behavioral outcomes.

##### 4.3.3.1. Effective information flow (EIF)

Among the simulation-derived metrics, those based on effective information flow (EIF) were consistently among the most predictive of aphasia severity across Sets 2b and 3. EIF quantifies the fidelity with which simulated signals propagate between regions within the modeled cortical network. Unlike functional connectivity-based measures, which infer relationships from correlated fluctuations, EIF provides a mechanistically grounded assessment of directed communication capacity by quantifying how effectively information propagates through a structurally perturbed system. These findings parallel empirical and computational studies showing that stroke disrupts directed information flow across large-scale networks, including reduced interhemispheric transfer of information, altered intrahemispheric coupling, and degraded information-processing capacity, all of which have been linked to motor, cognitive, and language impairments.^59,61–63^

An important observation is that regional EIF strength is not systematically reduced across the cortex following stroke (Figure 3); the average EIF strength in lesioned simulations remains nearly identical to the non-lesioned condition. This indicates that lesions do not globally diminish the system’s capacity for information transmission but instead redistribute communication pathways, degrading the flow from some regions while enhancing it for others. This phenomenon helps explain why EIF_RX_ strength similarity, a measure of how closely each region’s incoming information flow matches the healthy pattern, was consistently selected as a predictor in both Sets 2b and 3. Its importance suggests that changes in the spatial organization of directed communication, rather than uniform reductions in EIF magnitude, are most relevant for explaining aphasia severity. In parallel, several regional EIF strength metrics were also selected, indicating that localized increases or decreases in directed influence remain informative when interpreted within the broader context of disrupted network-level flow patterns.

The predictive importance of EIF provides strong support for the idea that aphasia severity reflects disruptions in how information is routed across large-scale networks, not simply where damage occurs – a pattern consistent with recent empirical and computational studies showing that stroke alters information flow and large-scale dynamical states in ways not captured by lesion characteristics alone. ^59,61–64^ Because EIF directly quantifies the capacity for signal transmission, it offers a mechanistically meaningful window into the functional consequences of structural lesions and is especially well suited to capturing chronic diaschisis, reduced integrative capacity, and breakdowns in hierarchical processing. These properties make EIF a promising tool not only for outcome prediction but also for testing mechanistic hypotheses about language network organization and, ultimately, for designing and evaluating targeted interventions that aim to restore or compensate for impaired information flow.

### 4.4. Future directions

The present findings highlight several avenues for extending and enhancing the REWIRED framework. First, although the current implementation relies solely on binary lesion maps, incorporating additional patient-specific biological markers, such as small vessel disease burden, cortical atrophy, white-matter hyperintensities, and metabolic or genetic risk factors, may further refine individualized predictions of language outcomes. Similarly, multimodal neuroimaging data, including diffusion tractography and resting-state or task-based fMRI, could be integrated with REWIRED-derived simulations to constrain or augment model estimates of structural and functional network integrity. Such hybrid approaches may offer complementary information, improving predictive accuracy beyond what either modeling or empirical imaging can achieve alone.

A second direction concerns the extension of REWIRED beyond static lesion effects to simulate longitudinal recovery trajectories. Future developments of the framework will incorporate biologically motivated plasticity rules that capture experience-dependent changes in connectivity and network organization following stroke. These recovery-focused simulations could enable in silico testing of rehabilitative interventions, including behavioral therapy dosing, task-specific training, or targeted neuromodulation strategies. Ultimately, dynamic modeling of both injury and recovery has the potential to support more precise, mechanism-informed prognostication and to guide individualized rehabilitation planning in clinical settings.

## 5. Limitations

Several limitations should be noted. First, the ARC dataset is predominantly composed of adults with chronic left-hemisphere stroke, which may limit the generalizability of the model to more diverse patient populations and to acute or subacute phases of recovery. Second, although the sample size is typical for lesion-symptom studies, it limits the precision of estimates and precludes robust subgroup or fairness analyses. The risk of overfitting was mitigated through rigorous nested cross-validation, which provides unbiased performance estimates, and the biological validity of the framework was further supported by empirical functional connectivity analyses performed in a separate CNLR cohort. Nevertheless, external validation of aphasia severity prediction performance in independent datasets will still be necessary to establish generalizability. Finally, because all predictors were derived from lesion masks and simulations without missing data, missingness did not introduce bias; however, future multimodal studies may require additional strategies for managing incomplete imaging or behavioral data.

## 6. Conclusion

This study provides support for the biological validity and predictive utility of the REWIRED framework and the parameterized mean field model on which it is based. Simulation-derived functional connectivity significantly predicted empirical functional connectivity beyond lesion burden and structural disconnection alone, indicating that the framework captures biologically meaningful aspects of large-scale network organization. Additionally, the strong aphasia prediction performance achieved using simulation-derived features, with simple binary lesion maps as the only subject-specific input, suggests that lesion-informed cortical dynamics simulations may provide a powerful and clinically scalable approach for linking structural injury to behavioral outcomes. Thus, the framework may provide a useful platform for investigating mechanistic hypotheses related to language function and recovery while improving the accuracy of clinical prognostication. By incorporating additional patient-specific data, such as small vessel disease burden, other neuroimaging markers, and metabolic and genetic factors, predictions are likely to improve further. Additionally, future extensions of the REWIRED framework will incorporate neuroplastic recovery simulations that could provide additional gains in prediction accuracy and serve as a clinical tool for guiding personalized rehabilitation strategies.

## 8. Declaration of generative AI and AI-assisted technologies in the manuscript preparation process

During the preparation of this work the author(s) used ChatGPT in order to assist with editing, revision, language refinement, and organizational restructuring of the manuscript. After using this tool/service, the author(s) reviewed and edited the content as needed and take(s) full responsibility for the content of the published article.

