## Supplementary Figure 1 for "Network measures from the REWIRED simulation framework enhance prediction of post-stroke aphasia severity"

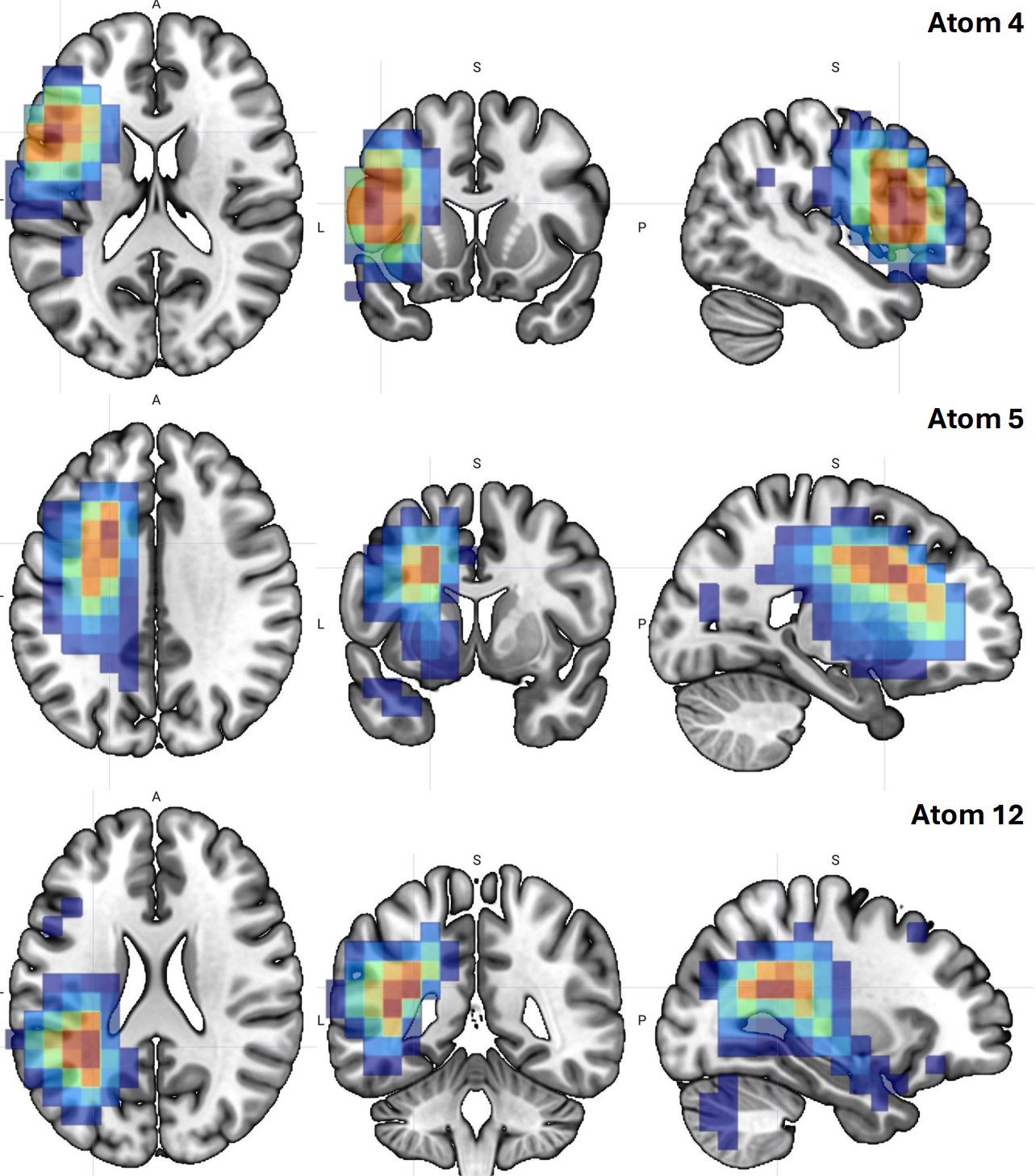

**Supplementary Figure 1**. Non-negative matrix factorization (NMF)–derived spatial lesion patterns (“atoms”). Each atom represents a recurrent multiregion lesion configuration identified across the cohort, shown in axial, coronal, and sagittal views. Warmer colors indicate regions with higher NMF loading (greater contribution to the atom). Atoms 4, 5, and 12 were consistently selected in models predicting aphasia severity. Atom 4 primarily captures inferior frontal and anterior perisylvian cortex involvement; Atom 5 reflects dorsal frontal and frontoparietal white-matter involvement; and Atom 12 reflects posterior temporal and temporoparietal white-matter involvement.
