## Supplementary Table 1 for "Network measures from the REWIRED simulation framework enhance prediction of post-stroke aphasia severity"

**Supplementary Table 1**. Global and regional mean field model parameters values as reported by Kong et al.^38^

| Global parameters |  |  |  |
| --- | --- | --- | --- |
| Synaptic coupling constant (*J*) | 0.2609 nA |  |  |
| Firing rate function slope (*a*) | 270 |  |  |
| Firing rate function threshold (*b*) | 108 Hz |  |  |
| firing rate function curvature (*d*) | 0.154 s |  |  |
| synaptic activity kinetics time constant (*τ_s_*) | 0.1 s |  |  |
| synaptic activity kinetics parameter (*r*) | 0.641 |  |  |
| **Regional parameters** | Recurrent connection strength (*w*) | External input current (*I*) | Noise scaling factor (*σ*) |
| *Left hemisphere* |  |  |  |
| Superior temporal sulcus (STS) | 0.494 | 0.313 | 0.00300 |
| Caudal anterior cingulate cortex (cACC) | 0.472 | 0.344 | 0.00201 |
| Caudal middle frontal gyrus (cMFG) | 0.818 | 0.278 | 0.00557 |
| Cuneus (CUN) | 0.150 | 0.329 | 0.00085 |
| Entorhinal cortex (EC) | 0.626 | 0.318 | 0.00350 |
| Fusiform gyrus (FG) | 0.370 | 0.333 | 0.00181 |
| Inferior parietal lobule (IPL) | 0.717 | 0.289 | 0.00476 |
| Inferior temporal gyrus (ITG) | 0.652 | 0.310 | 0.00384 |
| Isthmus of the cingulate cortex (ICC) | 0.657 | 0.263 | 0.00520 |
| Lateral occipital cortex (LOC) | 0.255 | 0.324 | 0.00151 |
| Lateral orbitofrontal cortex (lOFC) | 0.798 | 0.290 | 0.00512 |
| Lingual gyrus (LG) | 0.188 | 0.331 | 0.00099 |
| Medial orbitofrontal cortex (mOFC) | 0.919 | 0.280 | 0.00602 |
| Middle temporal gyrus (MTG) | 0.840 | 0.284 | 0.00551 |
| Parahippocampal gyrus (PHG) | 0.590 | 0.308 | 0.00361 |
| Paracentral lobule (PL) | 0.154 | 0.340 | 0.00055 |
| Inferior frontal gyrus pars opercularis (IFGop) | 0.655 | 0.302 | 0.00408 |
| Inferior frontal gyrus pars orbitalis (IFGorb) | 0.919 | 0.278 | 0.00608 |
| Inferior frontal gyrus pars triangularis (IFGtri) | 0.719 | 0.292 | 0.00470 |
| Pericalcarine cortex (PCAL) | 0.145 | 0.324 | 0.00097 |
| Postcentral gyrus (PostCG) | 0.156 | 0.341 | 0.00054 |
| Posterior cingulate cortex (PCC) | 0.589 | 0.305 | 0.00370 |
| Precentral gyrus (PreCG) | 0.208 | 0.332 | 0.00104 |
| Precuneus (PCUN) | 0.631 | 0.296 | 0.00413 |
| Rostral anterior cingulate cortex (rACC) | 0.937 | 0.284 | 0.00597 |
| Rostral middle frontal gyrus (rMFG) | 0.752 | 0.296 | 0.00475 |
| Superior frontal gyrus (SFG) | 0.720 | 0.300 | 0.00446 |
| Superior parietal lobule (SPL) | 0.316 | 0.331 | 0.00160 |
| Superior temporal gyrus (STG) | 0.374 | 0.331 | 0.00188 |
| Supramarginal gyrus (SMG) | 0.513 | 0.318 | 0.00294 |
| Frontal pole (FP) | 0.996 | 0.273 | 0.00658 |
| Temporal pole (TP) | 0.888 | 0.310 | 0.00499 |
| Transverse temporal gyrus (TTG) | 0.067 | 0.320 | 0.00070 |
| Insula (INS) | 0.402 | 0.340 | 0.00176 |
| *Right hemisphere* |  |  |  |
| Superior temporal sulcus (STS) | 0.392 | 0.320 | 0.00228 |
| Caudal anterior cingulate cortex (cACC) | 0.531 | 0.336 | 0.00253 |
| Caudal middle frontal gyrus (cMFG) | 0.744 | 0.287 | 0.00494 |
| Cuneus (CUN) | 0.161 | 0.330 | 0.00087 |
| Entorhinal cortex (EC) | 0.587 | 0.325 | 0.00310 |
| Fusiform gyrus (FG) | 0.337 | 0.335 | 0.00158 |
| Inferior parietal lobule (IPL) | 0.681 | 0.292 | 0.00451 |
| Inferior temporal gyrus (ITG) | 0.616 | 0.314 | 0.00355 |
| Isthmus of the cingulate cortex (ICC) | 0.607 | 0.268 | 0.00483 |
| Lateral occipital cortex (LOC) | 0.248 | 0.325 | 0.00144 |
| Lateral orbitofrontal cortex (lOFC) | 0.771 | 0.291 | 0.00496 |
| Lingual gyrus (LG) | 0.174 | 0.330 | 0.00095 |
| Medial orbitofrontal cortex (mOFC) | 0.908 | 0.282 | 0.00591 |
| Middle temporal gyrus (MTG) | 0.822 | 0.287 | 0.00532 |
| Parahippocampal gyrus (PHG) | 0.511 | 0.320 | 0.00288 |
| Paracentral lobule (PL) | 0.169 | 0.340 | 0.00064 |
| Inferior frontal gyrus pars opercularis (IFGop) | 0.577 | 0.314 | 0.00338 |
| Inferior frontal gyrus pars orbitalis (IFGorb) | 0.927 | 0.277 | 0.00613 |
| Inferior frontal gyrus pars triangularis (IFGtri) | 0.658 | 0.301 | 0.00412 |
| Pericalcarine cortex (PCAL) | 0.147 | 0.322 | 0.00104 |
| Postcentral gyrus (PostCG) | 0.154 | 0.341 | 0.00052 |
| Posterior cingulate cortex (PCC) | 0.585 | 0.304 | 0.00369 |
| Precentral gyrus (PreCG) | 0.183 | 0.335 | 0.00084 |
| Precuneus (PCUN) | 0.592 | 0.299 | 0.00387 |
| Rostral anterior cingulate cortex (rACC) | 0.918 | 0.288 | 0.00579 |
| Rostral middle frontal gyrus (rMFG) | 0.755 | 0.296 | 0.00476 |
| Superior frontal gyrus (SFG) | 0.715 | 0.302 | 0.00439 |
| Superior parietal lobule (SPL) | 0.303 | 0.334 | 0.00147 |
| Superior temporal gyrus (STG) | 0.356 | 0.336 | 0.00167 |
| Supramarginal gyrus (SMG) | 0.419 | 0.328 | 0.00221 |
| Frontal pole (FP) | 0.973 | 0.271 | 0.00652 |
| Temporal pole (TP) | 0.844 | 0.306 | 0.00492 |
| Transverse temporal gyrus (TTG) | 0.055 | 0.319 | 0.00068 |
| Insula (INS) | 0.376 | 0.340 | 0.00163 |
