## Supplementary Table 2 for "Network measures from the REWIRED simulation framework enhance prediction of post-stroke aphasia severity"

**Supplementary Table 2.** Complete list of features used in WAB AQ prediction pipeline

| **Feature class** | **Features** | **Feature set inclusion** |
| --- | --- | --- |
| Lesion distribution (n = 17) | Lesion volume  NMF atoms 1-16 | Set 1  Set 2a  Set 2b  Set 3 |
| Disconnectome (n = 70) | Global (n = 1)  Homotopic reserve*  Network-level (n = 1)  Left AF SC similarity  Regional (n = 68)  SC similarity  SC strength | Set 2a  Set 3 |
| Simulation (n = 146) | Global (n = 8)  FC similarity  FC mean  FC difference  FC variance  FC global efficiency  FC modularity  EIF_RX_ strength similarity  EIF_TX_ strength similarity  Network-level (n = 2)  FC FTP difference  FC FTP variance  Regional (n = 136)  FC strength  EIF_RX_ strength  EIF_TX_ strength  EIF local efficiency | Set 2b  Set 3 |
| **Abbreviations**: AF = arcuate fasciculus; FTPN = fronto-temporo-parietal network**; NMF = non-negative matrix factorization; SC = structural connectivity  *Homotopic reserve measures the extent to which the interhemispheric structural connectivity of regions homotopic to damaged left hemisphere regions is preserved.  ** Fronto-temporo-parietal network (FTPN) refers to a 12-region subset of left frontal, temporal, parietal, and peri-central cortex. Each metric was computed in the same manner as for the corresponding global feature but using the subgraph defined by these FTP regions, which included inferior parietal lobule (IPL), middle temporal gyrus (MTG), paracentral lobule (PL), inferior frontal gyrus pars opercularis (IFGop), IFG pars orbitalis (IFGorb), IFG pars triangularis (IFGtri), precentral gyrus (PreCG), rostral middle frontal gyrus (rMFG), superior frontal gyrus (SFG), superior temporal gyrus (STG), supramarginal gyrus (SMG), and transverse temporal gyrus (TTG). | | |
