## Supplementary Table 3 for "Network measures from the REWIRED simulation framework enhance prediction of post-stroke aphasia severity"

**Supplementary Table 3**. Selected features and model performances for all feature sets.

|  | **Tuned *k*** | **Selected Features*** | **# folds** | **RMSE**  **(mean ± s.d.)** | **MAE**  **(mean ± s.d.)** | **Pearson *r* (mean ± s.d.)** |
| --- | --- | --- | --- | --- | --- | --- |
| Set 1 | 8 | NMF atom 11  NMF atom 12  NMF atom 15  Lesion Volume  NMF atom 1  NMF atom 5  NMF atom 13  NMF atom 16 | 11  11  11  11  10  10  9  9 | 18.2 ± 3.9 | 14.5 ± 4.3 | 0.779 ± 0.088 |
| Set 2a | 7 | NMF atom 11  NMF atom 12  SMG SC similarity  Arcuate similarity  NMF atom 16  PHG SC similarity  TP SC strength | 11  11  11  10  8  8  8 | 18.4 ± 4.1 | 14.1 ± 4.6 | 0.772 ± 0.101 |
| Set 2b | 17 | FP EIF local efficiency  EIF_RX_ strength similarity  cMFG EIF_RX_ strength  ITG EIF_RX_ strength  mOFC EIF_TX_ strength  INS FC strength  FG FC strength  NMF atom 11  NMF atom 16  NMF atom 4  NMF atom 5  CUN EIF local efficiency  IFGop FC strength  NMF atom 12  NMF atom 14  PostCG EIF_RX_ strength  PCUN EIF_RX_ strength | 11  11  11  11  11  11  11  11  11  11  11  10  10  10  9  8  8 | 16.0 ± 2.5 | 12.4 ± 2.5 | 0.838 ± 0.077 |
| Set 3 | 16 | EIF_RX_ strength similarity  cMFG EIF_RX_ strength  ITG EIF_RX_ strength  mOFC EIF_TX_ strength  IFGop FC strength  FG FC strength  NMF atom 12  NMF atom 4  NMF atom 5  SMG SC similarity  rACC SC strength  FP EIF local efficiency  PCUN EIF_RX_ strength  rMFG EIF_RX_ strength  NMF atom 16  PostCG SC strength | 11  11  11  11  11  11  11  11  11  11  11  10  10  10  10  10 | 14.5 ± 3.3 | 10.9 ± 2.5 | 0.829 ± 0.127 |
| *****All region names refer to left hemisphere regions  **Abbreviations**: cMFG = caudal middle frontal gyrus; CUN = cuneus; EIF_RX_ = effective information flow (incoming); EIF_TX_ = effective information flow (outgoing); IFGop = inferior frontal gyrus pars opercularis; INS = insula; ITG = inferior temporal gyrus; FC = functional connectivity; FG = fusiform gyrus; FP = frontal pole; mOFC = medial orbitofrontal cortex; NMF = non-negative matrix factorization; PCUN = precuneus; PHG = parahippocampal gyrus; PostCG = postcentral gyrus; rACC = rostral anterior cingulate cortex; rMFG = rostral middle frontal gyrus; SC = structural connectivity; SMG = supramarginal gyrus; TP = temporal pole  **Definitions**: local efficiency = efficiency of communication among a region’s immediate neighbors; similarity = Pearson correlation with the non-lesioned counterpart of the same metric; strength = sum of connection weights for a region | | | | | | |
